# Variant-to-gene-mapping followed by cross-species genetic screening identifies GPI-anchor biosynthesis as novel regulator of sleep

**DOI:** 10.1101/2021.12.19.472248

**Authors:** Justin Palermo, Alessandra Chesi, Amber Zimmerman, Shilpa Sonti, Chiara Lasconi, Elizabeth B. Brown, James A. Pippin, Andrew D. Wells, Fusun Doldur-Balli, Diego R. Mazzotti, Allan I. Pack, Phillip R. Gehrman, Struan F.A. Grant, Alex C. Keene

**Affiliations:** Department of Biology, Texas A&M University, College Station, TX, United States; Center for Spatial and Functional Genomics, Children’s Hospital of Philadelphia, Philadelphia, PA, 19104, USA; Department of Pathology and Laboratory Medicine University of Pennsylvania Perelman School of Medicine Philadelphia PA USA; Division of Sleep Medicine/Department of Medicine, Perelman School of Medicine, University of Pennsylvania, PA, United States; Division of Medical Informatics, Department of Internal Medicine, University of Kansas Medical Center, Kansas City, KS, United States; Division of Pulmonary Critical Care and Sleep Medicine, Department of Internal Medicine, University of Kansas Medical Center, Kansas City, KS, United States; Department of Psychiatry, Perelman School of Medicine, University of Pennsylvania, PA, United States; Department of Pediatrics, The University of Pennsylvania Perelman School of Medicine, Philadelphia, PA, 19104, USA; Division of Human Genetics, Children’s Hospital of Philadelphia, Philadelphia, PA, 19104, USA; Department of Genetics, University of Pennsylvania, Philadelphia, PA, 19104, USA

**Author notes:** Denotes equal contributions (Justin Palermo, Alessandra Chesi, Amber Zimmerman). denotes contact PI.

## Abstract

Sleep is nearly ubiquitous throughout the animal kingdom, with deficiencies in sleep having been linked to a wide range of human disorders and diseases. While genome wide association studies (GWAS) in humans have identified loci robustly associated with several heritable diseases or traits, little is known about the functional roles of the underlying causal variants in regulating sleep duration or quality. We applied an ATAC-seq/promoter focused Capture C strategy in human iPSC-derived neural progenitors to carry out a ‘variant-to-gene’ mapping campaign that identified 88 candidate sleep effector genes connected to relevant GWAS signals. To functionally validate the role of the implicated effector genes in sleep regulation, we performed a neuron-specific RNAi screen in the fruit fly, *Drosophila melanogaster*. This approach identified a number of genes that regulated sleep, including phosphatidylinositol N-acetylglucosaminyltransferase subunit Q (*PIG-Q*), a gene that encodes an enzyme involved in the first step of glycosylphosphatidylinositol (GPI)- anchor biosynthesis. We show that flies deficient for *PIG-Q* have longer sleep during both day and night due to an increase in the total number of sleep bouts. Subsequent systematic investigation of other PIG-family genes identified increased sleep in flies for multiple different genes within the PIG pathway. We then mutated the *PIG-Q* locus in zebrafish and identified similar increases in sleep to those observed in *Drosophila*, confirming deep homology of *PIG-Q* mediated sleep regulation. These results provide the first physical variant-to-gene mapping of human sleep genes followed by a model organism-based prioritization, revealing a novel and conserved role for GPI-anchor biosynthesis in sleep regulation.

## INTRODUCTION

Dysregulation of sleep duration, timing and quality are associated with significant disease risk and public health burden ^1,2^. Sleep duration and quality vary dramatically between individuals, suggesting the presence of complex genetic factors that distinctly regulate characteristics of sleep ^3^. Despite this recognized concern, variable sleep differences across the population have a poorly understood biological basis, particularly from the genetic standpoint ^4^.

Virtually all physiologic processes are impacted by sleep, strongly suggesting that its function extends beyond the brain to affect diverse cell types and physiological processes ^4^. In recent decades, significant progress has been made in understanding the mechanisms underlying circadian rhythms, including the identification of many genetic loci that impact inter-individual variability, yet, much less is known about variability in sleep disorders such as insomnia across human populations ^5^. A number of GWAS efforts have been conducted for insomnia -related phenotypes. Initial efforts in relatively smaller datasets (N≤10,000) failed to achieve genome-wide significant associations with self-reported insomnia symptoms ^6,7^. However, more recent studies have combined data from the UK Biobank and 23andMe for an insomnia GWAS of >1.3 million individuals that yielded 202 associated loci significant at the genome-wide level ^8^.

A central impediment to interpreting GWAS studies for complex traits is determining whether the nearest gene to an associated SNP functionally contributes to the observed phenotype ^9,10^. Even when the most obvious gene at the locus would appear functionally linked a *priori*, perhaps those genes represent a ‘red herring’ and the actual causative gene remains to be discovered, or equally likely, there may be more than one effector gene at a given locus ^11–13^. While there is a relative paucity of public domain genomic data relevant to sleep-related tissue, such as eQTL data, related techniques can be leveraged to identify sleep influencing effector genes. We elected to carry forward established and novel insomnia GWAS signals to such a next level of investigation. The application of Chromatin Conformation Capture based ultra-high resolution promoter ‘interactome’ have the ability to determine if chromatin ‘looping’ contributes to human disease at key locations associated with complex traits ^14–18^.

Given the need for functional insight into reproducible genetic associations with sleep traits, our goal was to provide the first comprehensive physical variant-to gene-mapping for insomnia GWAS-implicated loci by taking advantage of our data generated on neural progenitor cells (NPCs). The leveraging of GWAS findings to discover genetic variation that impacts sleep require first defining the effector genes impacted by the key regulatory regions harboring the associated putative causal non-coding SNPs, then localizing expression to defined brain regions or cell types, and finally, characterization of impact on sleep duration and timing *in vivo*. Here, we integrated ATAC-seq/promoter-focused Capture C data with GWAS findings to implicate effector genes impacted by regulatory regions coinciding with key insomnia-associated SNPs with cell-type specificity. These data provide a list of candidate sleep regulators, and provide the basis for *in vivo* analysis of gene function in genetically amenable model systems. We first used a genome-wide RNAi library in the fruit fly, *Drosophila melanogaster*, to test whether the candidate genes function in neurons to regulate sleep. These experiments were followed by CRISPR-based mutagenesis in zebrafish to determine whether the effects identified in flies are conserved in a vertebrate model. These efforts identified numerous novel regulators of sleep, including a role for *PIG-Q* and GPI-anchoring in sleep regulation. Furthermore, this integrative approach provides a framework for high-throughput validation of candidate genes implicated through the integration of GWAS signals with variant-to-gene mapping in a relevant human cell model followed by *in vivo* phenotypic analyses in animal models.

## RESULTS

To search for potential regulators of human sleep, we first leveraged genome-wide significant signals from published insomnia GWAS, derived from a combination of the UK Biobank cohort and individuals who were genotyped by 23andMe and consented to participate in research for insomnia^19^. A total of 202 genome-wide significant loci previously implicated 956 genes through positional, eQTL and chromatin mapping that were enriched for neural cell types. These genetic associations provided the basis for variant-to-gene mapping and functional validation of sleep genes **(Fig 1a)**.

**Figure 1.**
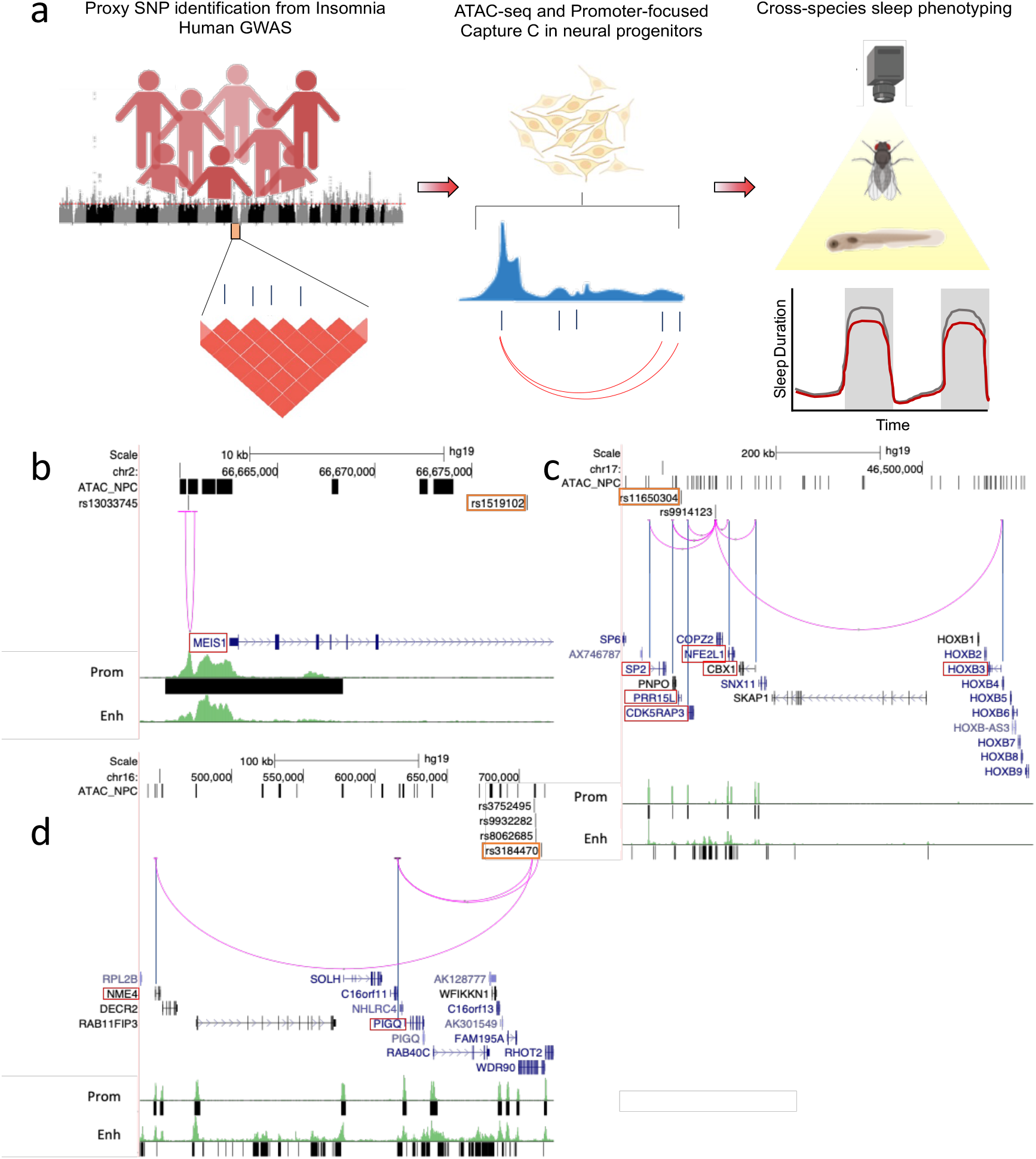
Translating human GWAS signals to functional outcomes with variant-to-gene mapping. **a**, Leveraging existing insomnia human GWAS loci, we identified proxy SNPs in strong linkage disequilibrium with sentinel SNPs using both genome wide ATAC-seq and high-resolution promoter-focused Capture C data from iPSC-derived neural progenitor cells then performed high-throughput sleep and activity screening using *Drosophila* RNAi lines with confirmation in a vertebrate zebrafish (*Danio rerio*) model. **b-d**. Three examples of chromatin loops linking insomnia associate SNPs to candidate effector genes in neural progenitor cells. **b**, rs13033745 (r^2^ with sentinel SNP rs1519102 = 0.84) loops to the *MEIS1* promoter region. **c**, rs9914123 (r^2^ with sentinel SNP rs11650304 = 0.76) loops to the promoters of *SP2, PRR15L, CDK5RAP3, NFE2L1, CBX1* and *HOXB3* in a ∼700 kb region. **d**, rs3752495, rs8062685 and rs9932282 (r^2^ with sentinel SNP rs3184470 = ∼1) loop to the promoters of *PIG-Q, NHLRC4* and *NME4*. Orange box: sentinel SNP. Black bars: open chromatin peaks from ATAC-seq. Magenta arcs: chromatin loops from promoter-focused Capture C. Neuronal enhancer and promoter tracks are from ^79^.

While sleep impacts tissues throughout the body, it is largely defined by physiological changes in brain activity that drive sleep-associated behaviors ^20,21^. To examine the effects of loci identified through GWAS for insomnia in brain-related cell types, we leveraged data derived from both genome wide ATAC-seq and high-resolution promoter-focused Capture C data in order to implicate insomnia effector genes contacted directly by regulatory regions harboring the GWAS-associated variants. Since both neurons and glia regulate many aspects of sleep function, we focused our initial analysis on iPSC-derived neural progenitor cells, which are the precursors from which most of the glial and neuronal cell types of the CNS originate ^22–24^. iPSCs derived from two healthy individuals (CHOPWT10 and CHOPWT14) were differentiated to NPC and cultured using standard techniques ^22^. We employed a high-resolution genome-scale, promoter-focused Capture C-based approach^14^ that utilizes a 4-cutter restriction enzyme (DpnII, mean fragment size 433 bp; median 264 bp), achieving higher resolution than the more commonly used 6-cutter Hi-C related approaches (HindIII, mean fragment size 3,697 bp; median 2,274 bp) ^14,25^.

We leveraged Capture C and ATAC-seq data generated from the same cell lines and sequenced on the Illumina platform ^22^. The ATAC-seq data was analyzed with the ENCODE ATAC-seq pipeline (https://github.com/kundajelab/atac_dnase_pipelines) with the “optimal” IDR calling strategy, yielding 100,067 open chromatin peaks. To determine the informative genetic variants associated with insomnia, we extracted 11,348 proxy SNPs for each of 246 independent signals coinciding with 200 informative insomnia GWAS loci where proxies could be identified (r^2^>0.7 to sentinel SNP in Europeans) and overlapped those variants with the positions of the open chromatin regions (ATAC-seq peaks). We identified 321 informative proxy SNPs corresponding to 100 of the insomnia loci in high linkage disequilibrium (LD) with the sentinel SNP at each locus investigated. This effort substantially shortened the list of candidate causal variants from the initial GWAS discoveries, since ATAC-seq permitted us to focus on variants residing within open chromatin regions in cells that are relevant for sleep-wake regulation.

Leveraging the Capture C dataset ^22^, we mapped the informative variants from the insomnia GWAS loci to their target genes in NPCs. Of the insomnia GWAS loci investigated, 36 were implicated in a chromatin loop, with proxy SNPs residing in open chromatin (not in a baited promoter region) contacting one or more open gene promoters. A total of 135 open baited regions corresponding to the promoters of 141 genes (88 coding) were connected to 76 open chromatin regions harboring one or more insomnia proxy SNP through 148 distinct non bait-to-bait chromatin looping interactions (**Table S1**). Some chromatin loops pointed to the nearest gene (such as rs13033745 at *MEIS1*; **Fig 1b**), while others to a gene or multiple genes further away from the candidate regulatory open SNP (such as rs9914123, which resides in an intron of *COPZ2* but loops to the promoters of several genes in a ∼700 kb region; **Fig 1c**). The chromatin loops involving 3 insomnia-associated SNPs (rs3752495, rs8062685 and rs9932282; r^2^ with sentinel SNP rs3184470 = ∼1) and the promoters of *PIG-Q* and *NHLRC4* and *NME4*are shown in **Fig 1d**. These analyses identified 88 candidate target coding genes, including *MEIS1* which has already been widely implicated in sleep and restless leg syndrome^26,27^. Mining our RNA-seq data on the same cell line, we observed that almost all of the identified target coding genes (80/88) were expressed at moderate or high level (percentile of expression >50%, TPM > 1.5) (**Table S1**).

The genes and neural mechanisms regulating sleep are highly conserved from flies to mammals, and powerful genetics in non-mammalian models can be used to screen for novel regulators of sleep ^28,29^. In fruit flies, sleep can be identified through behavioral inactivity bouts lasting for 5 minutes or longer, and the *Drosophila* Activity Monitor (DAM) System detects activity through infrared beam crossing and is widely used to quantify sleep ^30,31^. To determine whether the candidate genes from our 3D genomics effort contribute to sleep regulation, we expressed RNAi targeted in candidate genes selectively in neurons under control of nSyb-GAL4, and screened genes for sleep (**Fig 2a**). Of the 88 insomnia-associated coding genes identified through our variant-to-gene mapping analyses, we could identify 66 genes with moderate to strong orthologs in fruit flies. Of these genes, 54 had available RNAi lines in the Vienna Drosophila Stock Center (VDRC) or the DRSC/TRiP collection (**Fig 2a**). There were no significant differences in total sleep duration between control flies from each RNAi library and, therefore, all lines were tested and analyzed together (**Fig 2b**). This initial analysis identified a number of short and long sleeping lines. For example, knockdown of the genes encoding the cell adhension molecule *connectin*, (ortholog of CHADL) and the BHLH transcription factor, *daughterless* (ortholog of TCF12), resulted in short sleeping phenotypes (**Fig 2b, Table S2**). In addition, we identified a short-sleeping phenotype for the Hox cofactor, *homothorax* (*hth*), and ortholog of mammalian *Meis1* which has already been implicated in sleep and human restless leg syndrome (Sarayloo et al., 2019). The screen also identified a number of genes associated with long-sleeping phenotypes including *Gß13F* (ortholog of GNB3), the RNA helicase *twister* (ortholog of SKIV2L), and the glycosylphosphatidylinositol anchoring biosynthesis protein, *PIG-Q* ^33^. *PIG-Q* encodes an enzyme N-acetylglucosaminyl transferase that localizes to the endoplasmic reticulum and is required for synthesis of the GPI anchor that regulates the cellular localization of ∼150 proteins. Together, these findings revealed complex sleep phenotypes associated with individual genes identified through human GWAS studies. Given the high conservation of *PIG-Q* and its critical role in GPI biosynthesis, we chose to focus on this gene for further analyses.

**Figure 2.**
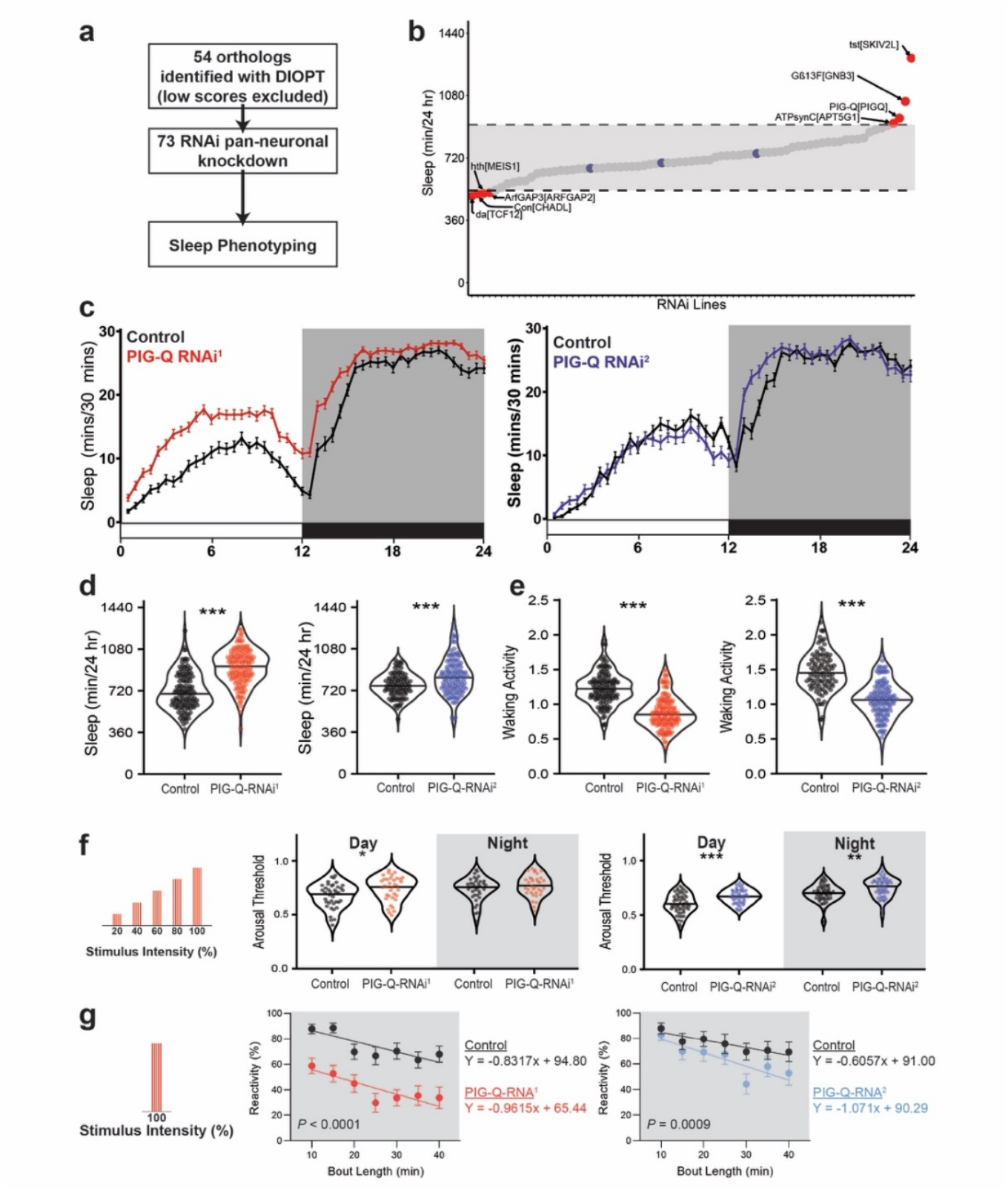
*PIG-Q* knockdown increases sleep duration and sleep depth. **a**, Design of orthologous gene screen. **b**, Total sleep minutes over a 24-h period in viable RNAi crosses (73 lines, n > 16 per line). Dotted lines and greyed area indicate two std. dev from the mean for every animal tested in either direction. Blue dots indicate control sleep responses, while red dots indicate sleep responses of RNAi lines that fall outside two std. dev. **c**, Sleep profiles of two independent RNAi lines targeting *PIG-Q* (*PIG-Q*-RNAi^1^ : red; *PIG-Q*-RNAi^2^ : blue). **d**, Knockdown of *PIG-Q* in significantly increases total sleep (t-test, *PIG-Q*-RNAi^1^ t_260_ = 11.42; *P*<0.0001; *PIG-Q*-RNAi^2^ t_184_ = 4.282; *P* = <0.0001). **e**, Knockdown of *PIG-Q* in significantly decreases waking activity (t-test, *PIG-Q*-RNAi_1_ t_260_ = 11.45; *P* = <0.0001; *PIG-Q*-RNAi_2_ t_184_ = 11.09; *P*<0.0001). (**f-h**), The Drosophila ARousal Tracking (DART) system records fly movement while simultaneously controlling periodic mechanical stimuli. **f**, The DART system was used to measure arousal threshold and reactivity. Arousal threshold was measured on sleeping flies using mechanical stimuli of increasing strength. Reactivity was measured by assessing the proportion of flies that react to a single mechanical stimulus for each bin of immobility. **g**, Knockdown of *PIG-Q* significantly increases arousal threshold (REML, *PIG-Q*-RNAi ^1^: F_1,73_ = 4.267, *P*=0.0424; *PIG-Q*-RNAi^2^ : F_1,102_ = 16.42, *P*<0.0001). This occurs during the day for both independent RNAi lines (*PIG-Q*-RNAi^1^ *P*=0.0127; *PIG-Q*-RNAi^2^ : *P*=0.0002), while an increase in arousal threshold only occurred in one line during the night (*PIG-Q*-RNAi^1^: *P*=0.4308; *PIG-Q*-RNAi^2^ : *P*=0.0020). **h**, Knockdown of *PIG-Q* significantly decreases nighttime reactivity (ANCOVA with bout length as covariate, *PIG-Q*-RNAi^1^: F_1,661_ = 107.1, *P*<0.0001; *PIG-Q*-RNAi^2^: F_1,594_ = 24.87, *P*<0.0001). For sleep profiles, error bars represent +/-standard error from the mean. For violin plots, the median (solid black line) is shown. White background indicates daytime, while gray background indicates nighttime. \**P*<0.05, \*\**P*<0.01, \*\*\**P*<0.001, \*\*\*\**P*<0.0001.

To validate the screening results, we repeated the sleep analyses using additional genetic controls including a second, independently-derived RNAi line. Flies with pan-neuronal *PIG-Q* were compared to controls harboring the GAL4 driver or the RNAi line alone. Both RNAi lines significantly increased sleep over control flies, fortifying the notion that loss of *PIG-Q* in neurons promotes sleep during the daytime and nighttime **(Fig 2c-d, Fig S1)**. The waking activity, defined as the average amount of activity while the animal is awake, was reduced in *PIG-Q* knockdown flies, suggesting a role in activity in addition to sleep regulation (**Fig 2e**). Further, the identified phenotypes were present in male flies, revealing that the effect of *PIG-Q* knockdown on sleep is not sex-specific **(Fig S2)**. Together, these results confirmed knockdown of *PIG-Q* in neurons promotes sleep.

Across phyla, sleep is defined by a homeostatic rebound following deprivation and reduced responsiveness to external stimuli. To determine whether sleep homeostasis is disrupted in *PIG-Q* knockdown flies, we sleep-deprived flies for 12-hours during the lights off period (ZT12-24) by mechanical stimulation and measured recovery sleep. Following deprivation, both control and *PIG-Q* knockdown flies significantly increased sleep. A direct comparison revealed a similar percent increase in flies suggesting homeostatic rebound is intact in *PIG-Q* deficient flies **(Fig S3**). To further investigate the role of *PIG-Q* in sleep regulation we quantified arousal threshold in knockdown flies, using the Drosophila Arousal Threshold (DART) system ^34,35^. Analysis of video recordings in this system confirmed the increased sleep phenotype of *PIG-Q* knockdown flies from infrared tracking **(Fig S4)**. To probe for sleep depth, sleep was recorded and analyzed by video-tracking prior to and following exposure to mechanical shaking that increase in strength. There was an increase in daytime arousal threshold in flies with pan-neuronal expression of either *PIG-Q* RNAi line, suggesting loss of *PIG-Q* increases sleep depth. Nighttime reactivity was reduced in both lines, suggesting a role for *PIG-Q* in sleep depth **(Fig 2e)**. No differences in reactivity were identified during the daytime in *PIG-Q* knockdown flies, suggesting *PIG-Q* differentially effects sleep depth during the day and night. **(Fig S5)**. To confirm these results, we analyzed arousability as a function of sleep bout length. Awakenings induced by mechanical stimulus was reduced during minutes 10-40 of sleep in flies with pan-neuronal knockdown of *PIG-Q* **(Fig 2f)**. These results strengthen the finding that knockdown of *PIG-Q* expression increases sleep depth, particularly during longer sleep bouts.

In *Drosophila* and mammals, sleep regulating neurons are found in numerous brain regions ^23,28^. To localize the effects of *PIG-Q* we selectively knocked down function in different populations of neurons within the brain and measured the effects on sleep. There was no effect of knockdown in a number of canonical sleep areas including the mushroom body (R69F08) and the c929-driver that labels numerous sleep-regulating peptidergic neurons (Park et al., 2008). Therefore, *Pig-Q* is unlikely to generally impact cellular function within sleep regulating circuits. However, knockdown in cholinergic neurons within the brain (Cha-GAL4) significantly increased sleep, phenocopying pan-neuronal knockdown (**Fig 3a**), suggesting that *PIG-Q* modulates the function of these excitatory neurons. We also found increased sleep when *PIG-Q* was knocked down in a number of neuronal types including the circadian pacemaker PDF neurons, 5**-**HTR neurons, the ellipsoid body and fan-shaped body (**Fig 3a; Table S3)**. Importantly, two drivers that label the sLNvs pacemakers cells (PDF-GAL4 and dpp-GAL4) both increase sleep^37,38^. These findings suggest *PIG-Q* is required in diverse subsets of sleep regulating neurons for normal sleep. Therefore, *PIG-Q* is likely to function in multiple subsets of neuromodulatory circuits to regulate sleep.*PIG-Q* functions in the glycosylphosphatidyinositol (GPI) biosynthesis pathway that is highly conserved and critical for the function of GPI-anchored proteins (Kinoshita, 2020). Given the role of GPI-anchored proteins in sleep regulation ^40^, we sought to determine whether additional components of this pathway are involved in sleep regulation. In total we tested all 18 genes associated in the PIG-associated biosynthesis pathway (**Fig 3b)**. We knocked down 18 genes individually in different experiments in the GPI-biosynthesis pathway pan-neuronally and measured the effect on sleep. Sleep was significantly increased in flies with loss of PIG-Z, PIG-L, PIG-O, PIG-C, PIG-G, and PIG-M, where all slept longer than control flies expressing the RNAi line alone or the Nsyb-GAL4 driver alone (**Fig 3c, Table S4)**. The majority of genes targeted for pan-neuronal knockdown in this pathway resulted in increased sleep. Together, these findings suggest that generalized disruption of PIG-mediated GPI-biosynthesis promotes sleep.

**Figure 3.**
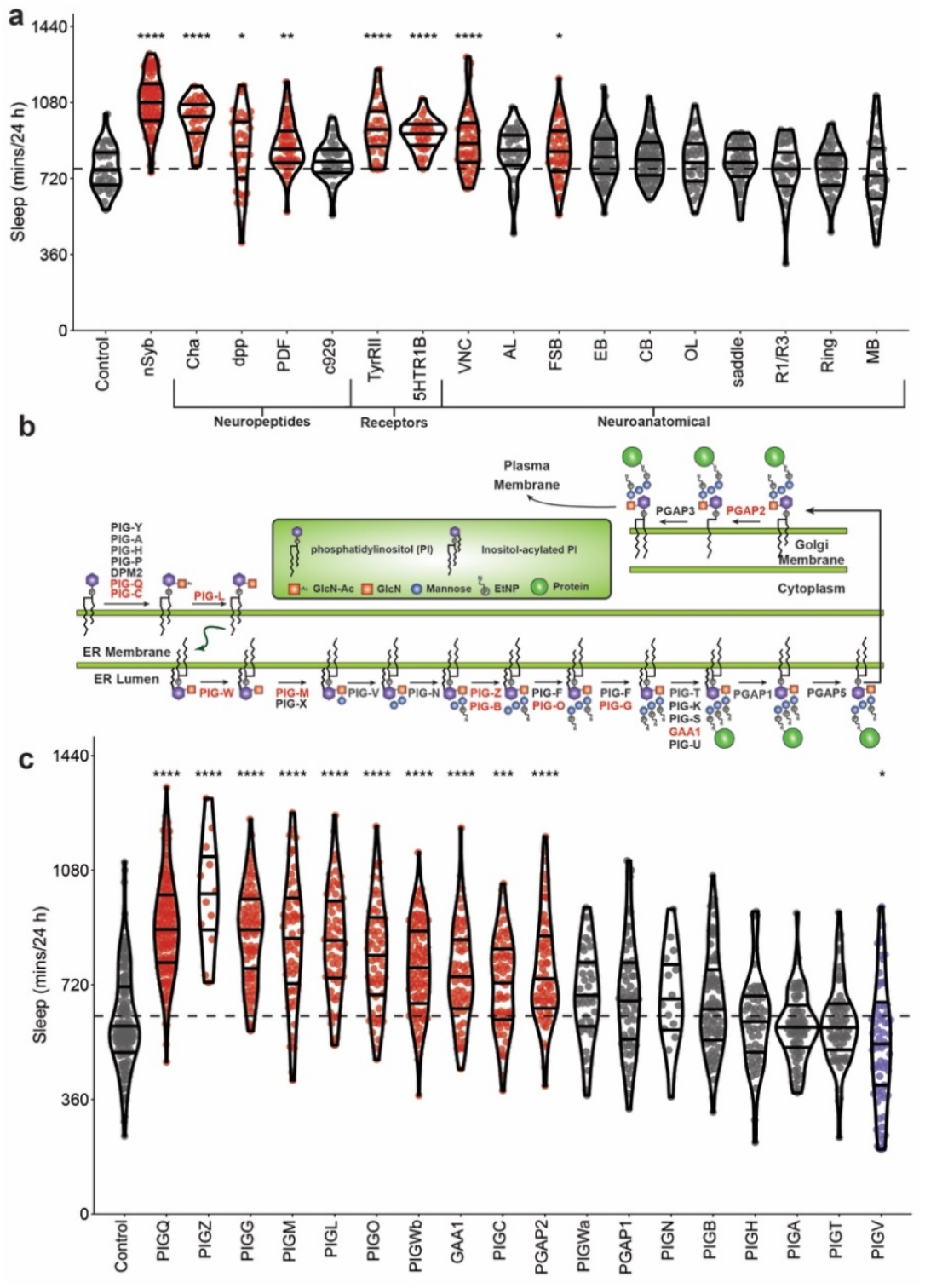
Localization of *PIG-Q* and characterization of the GPI-anchor biosynthesis genes in sleep regulation. **a**, Knockdown of *PIG-Q* in multiple *Drosophila* neuronal subpopulations affects sleep duration (ANOVA, F_18,756_ = 25.21, *P*<0.0001). The dotted line represents the mean of the control line. **b**, Phosphatidylinositol glycan anchor biosynthesis (PIG) pathway. Genes highlighted in red represent genes that show long sleep phenotypes when knocked down pan-neuronally in Drosophila as described below, while genes in grey exhibited no or short sleep phenotype. Genes in black were untested because there were no available RNAi lines. **c**, Knockdown of multiple genes in the PIG pathway affects sleep duration (ANOVA, F_18,1251_ = 39.63, *P*<0.0001). The dotted line indicates the mean of the control line. For violin plots, the median as well as 25^th^ and 75^th^ percentiles are shown (solid black lines). Each dot represents an individual fly; red indicates sleep duration that is significantly higher than the control, gray indicates sleep that is not significantly different, while blue indicates sleep that is significantly lower than the control as revealed by Dunnett’s multiple comparisons test. \**P*<0.05, \*\**P*<0.01, \*\*\**P*<0.001, \*\*\*\**P*<0.0001.

*PIG-Q* is a conserved gene across species with 44% amino acid sequence similarity between *Drosophila* and humans ^41^. Conservation is higher among vertebrates with a sequence similarity of 77% between zebrafish and humans (Hu et al., 2017). To determine if the functional effects of sleep in *Drosophila* are conserved in vertebrates, we examined the role of *PIG-Q* on sleep in zebrafish, a leading vertebrate model of sleep ^42^. We disrupted *PIG-Q* expression using CRISPR/Cas9 gene editing in zebrafish (*Danio rerio*). Targeted biallelic genetic mutations producing high-efficiency knockouts (KO) were generated in F0 larvae ^43^ (**Fig 4b-c**). We selectively targeted exon 7, which is conserved across both zebrafish *PIG-Q* transcripts and is a highly conserved region across species ^44^ (**Fig 4a**). Exon 7 is also part of the N-acetylglucosaminyl transferase component, a major functional component of the PIG-Q protein ^45^. Five days post fertilization (dpf), *PIG-Q* KO larvae were screened for sleep phenotypes compared to control zebrafish (scrambled gRNA-injected) larvae (**Fig 4a**). Behavioral analyses were performed using standardized methodology in zebrafish that has been previously used for genetic and pharmacological screens ^46,47^. The fish were genotyped immediately following behavioral analysis, which confirmed a mutation efficiency criterion for inclusion of >90% (**Fig 4c**). As with *Drosophila*, loss of *PIG-Q* significantly increased sleep duration during the night (*P*<0.001) compared to scrambled gRNA-injected controls (**Fig 4h-i**). Daytime sleep duration was also significantly (p<0.05) increased compared to controls (**Fig 4e and h**) with an increase in sleep bout number (p<0.05) (**Fig 4g**). The sleep differences at night were due to increased sleep bout length (**Fig 4k**) rather than sleep bout number (**Fig 4j**). This further supports the notion that loss of *PIG-Q* function increases sleep consolidation at night. However, there was not a significant change in overall activity during the day (**Fig 4d and f**), indicating that in zebrafish, *PIG-Q* exerts its effects primarily on sleep regulation rather than locomotion. Taken together, these findings confirm that loss of *PIG-Q* increases sleep across phylogeny.

**Figure 4.**
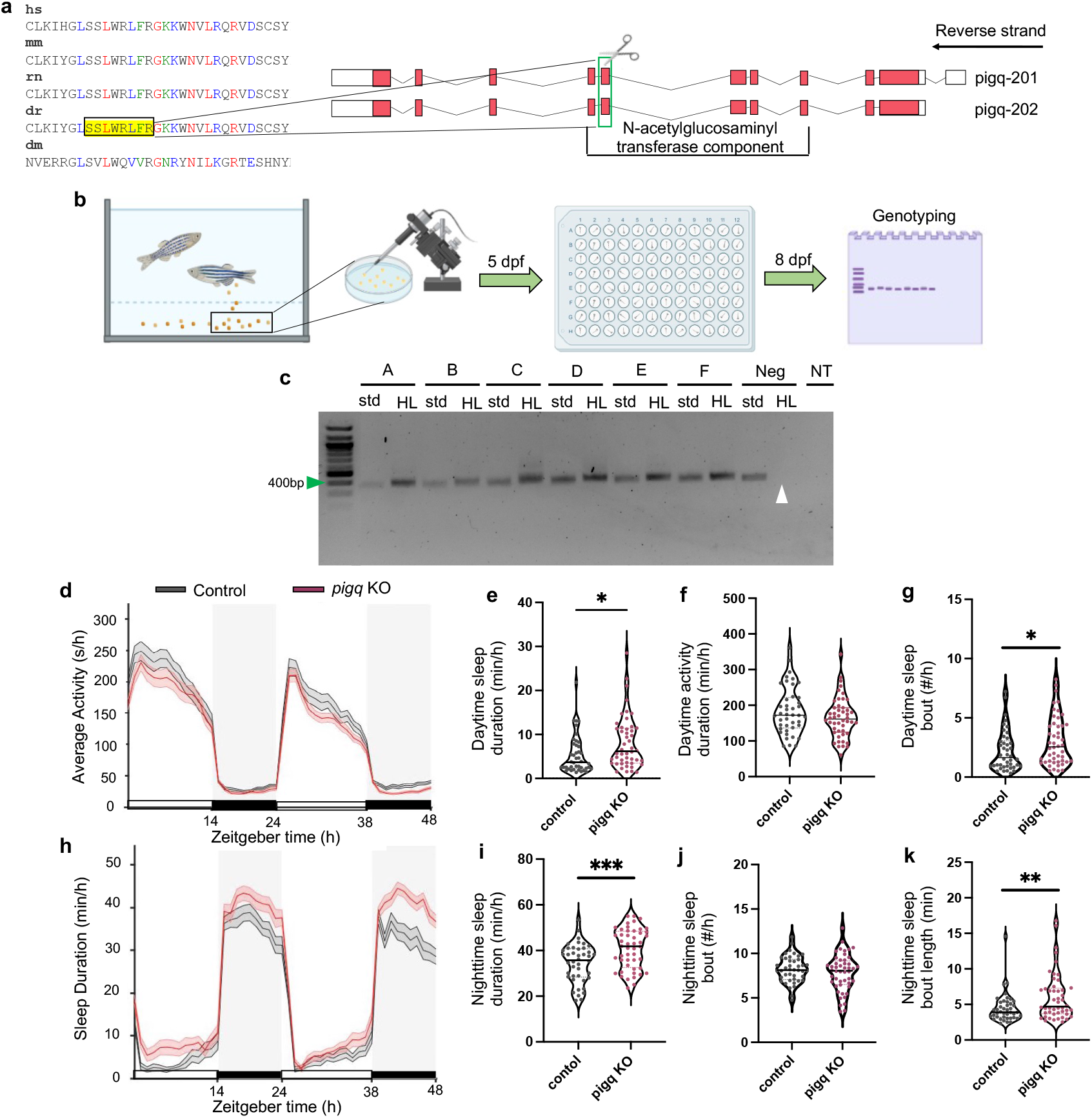
CRISPR mutation of *PIG-Q* in zebrafish increases sleep. **a**, CRISPR sgRNA design **b**, Schematic of experimental design from embryo injection to CRISPR mutation confirmation. **c**, Representative gel used for genotyping. Green arrow indicates 400bp on the ladder. Expected PCR product was 366bp. White arrow indicates wild-type DNA suppression using HL PCR as a negative control. NT is no template. **d**, Average (± s.e.m.) activity for 48 hours beginning at lights on (9am). **e**, Cumulative daytime sleep across both light periods (9am-11pm) was increased in *PIG-Q* KOs (mean diff: 2.83 ± 1.09, t(85.2) = 2.59, *P* = 0.04). **f**, No difference was found in daytime activity (mean diff: -19.3 ± 13.01, t(88) = 1.48, *P* = 0.14). **g**, Daytime sleep bout number was increased in *PIG-Q* KOs (mean diff: 0.83 ± 0.40, t(88) = 2.06, p = 0.04). **h**, Average (± s.e.m.) sleep duration across 48 hours beginning at lights on (9am). **i**, Cumulative nighttime sleep duration was increased in *PIG-Q* KOs across both dark periods (11pm-9am) (mean diff: 6.38 ± 1.82, t(88) = 3.5, *P* = 0.0007). **j-k**, Nighttime sleep bout number did not differ between groups (**j**, mean diff: -0.41 ± 0.38, t(85.2) = 1.08, *P* = 0.28), but nighttime sleep bout length was increased in *PIG-Q* KOs (**k**, mean diff: 1.60 ± 0.58, t(79.7) = 2.75, *P* = 0.007). Animals were kept on a 14:10 light:dark cycle. Gray shaded boxes indicate night, while white represents day. N = 42 scramble-injected controls, 48 *PIG-Q* KOs. Independent student’s t-test was used to compare *PIG-Q* KOs and controls. Welch’s correction was applied to e, j, and k because unequal variances between groups were determined. \**P*<0.05, \*\**P*<0.01, \*\*\**P*<0.001.

## DISCUSSION

We have conducted the first physical variant-to-gene mapping for insomnia GWAS by identifying putative causal variants and their associated effector genes leveraging data from an ATAC-seq/chromatin conformation capture-based approach, followed by assessing functional effects on sleep/wake regulation in *Drosophila* and zebrafish. The detailed behavioral platforms to characterize sleep duration and intensity, availability of RNAi libraries that allow for genome-wide *in vivo* analysis of sleep function, and high throughput assays make *Drosophila* an excellent system for validating the role of putative regulators of sleep ^23,29^. The candidate genes derived from our analysis were subjected to a neuron-specific RNAi screen in *Drosophila melanogaster* followed by in-depth sleep phenotyping in *Drosophila* and zebrafish to assess the impact of such genetic perturbation on sleep/wake regulation. As a consequence, a number of short- and long-sleeping lines were identified. Therefore, this approach provides proof-of-principle for the use of genetic models to interrogate the functional roles of genes implicated through human GWAS studies on complex behavior.

Screening identified multiple genes with short or long sleep phenotypes including numerous transcription factors. These genes provide candidates for further validation in flies, including verifying phenotypes in classic genetic mutants and localizing the effects of the genes. We focused functional validation on the *PIG-Q* gene because of the robustness of the phenotype and a previously identified role for GPI-anchored genes in sleep regulation ^40^. Knockdown of *PIG-Q* significantly increased both daytime and nighttime sleep. Restricting knockdown to a number of different neuromodulatory neurons, including cholinergic and tyraminergic neurons that have previously been implicated in sleep^48,49^ phenocopying pan-neuronal knockdown, supporting the notion that *PIG-Q* modulates sleep through its effect in these neuronal groups. To examine how *PIG-Q* regulates sleep we examined sleep and circadian regulation across a number of contexts. We subsequently localized *PIG-Q* function to numerous populations of neurons including, cholinergic neurons, tyraminergic neurons, and neurons expressing the serotonin receptor 5HT1B, revealing *PIG-Q* is likely to act in broad classes of neurons to regulate sleep. The sleep phenotype was subsequently recapitulated in the zebrafish model demonstrating conservation of *PIG-Q* function in regards to sleep function. The identification of *PIG-Q* implicates GPI-linked proteins in sleep regulation. Mutations in the GPI-anchored cell-surface protein *sleepless* leads to robust reductions in sleep ^40,50^. In total, the *Drosophila* genome encodes ∼150 GPI linked proteins^51,52^, and systematically testing the role of these in sleep regulation may uncover genes that are downstream of the GPI biosynthesis pathway and a broader role for GPI anchoring in sleep regulation.

In line with our similar work in other traits ^14,17,18,22^, we applied a physical variant-to-gene mapping approach to identify candidate regulators of sleep, using loci derived from GWAS studies. While a number of studies have used human GWAS to develop candidate regulators of sleep that can be used for genetic screening, this approach may lead to the incorrect genes being implicated. For example, GWAS efforts by others for obesity have clearly shown a pronounced association with variation within the *FTO* gene that associate with obesity ^53^. This robust association signal resides within an intronic region of this gene ^53^ and has gone on to be widely replicated in other ethnicities ^54–56^, plus children ^57^. Although many publications have now studied the role of the *FTO* locus in the context of obesity, a number of studies^46-48^ demonstrated that *FTO* is in fact likely NOT the principal causal effector gene for obesity at this locus, but rather it is *IRX3* and *IRX5* ^11–13^, suggesting that the genetic variant resides in an enhancer embedded in one gene that influences the expression of others. As such, despite a great deal of data implicating *FTO* as the gene involved in obesity, in fact through refined methodologies (similar to what we propose herein) in the absence of eQTL support, other genes that are physically located near *FTO* are actually the physiologically relevant effector genes ^11–13^. Similarly, the insomnia-associated candidate regulatory variants at GWAS locus number 170 identified by our variant-to-gene mapping reside in an intron of *WDR90*, but loop across ∼90 kb to the promoter region of two candidate effector genes farther away, *NHLRC4* and *PIG-Q*, and to the *NME4* promoter, 258kb away. While further experiments, such as CRISPR/Cas9 editing of the candidate variants in human cells, are required to validate a regulatory role on these target genes, our *Drosophila* phenotypic screen identified *PIG-Q* as the likely culprit gene at this locus.

Large-scale genetic screens have been applied in a number of animal models including *C. elegans, Drosophila*, zebrafish and mice to identify genetic regulators of sleep ^40,46,58–61^. Despite the ability for high-throughput behavioral screening, surprisingly few studies have used these models to validate genes identified in human GWAS studies. In fruit flies, the voltage gated Ca^2+^ channel cacophony ^62^ and the K(ATP) channel ABCC9 ^63^ have been identified as sleep regulators following their identification in human GWAS. In addition, cross-species analysis has found the *epidermal growth factor receptor* (EGFR) promotes sleep in *C. elegans, Drosophila*, and zebrafish, and is associated with variation in human sleep duration and quality ^64–66^. Similar approaches used in NPCs, can be applied to broader cell types including glia, insulin regulating cells, and the fat body (adipose tissue) all of which have been found to be regulators of sleep ^23,28^. Therefore, the application of model organisms combined with variant-to-gene mapping has potential to identify novel genetic regulators for many traits that have been studied using GWAS.

Taken together, this study provides a proof-of-principle application of physical gene-variant mapping and screen-based *in vivo* validation for a complex behavior. This approach identified PIG family proteins as conserved regulators of sleep, and raises the possibility that differences in GPI-biosynthesis contribute to naturally occurring variation in sleep. This study also provides a framework for interrogation of the large number of results emerging from other GWAS of sleep and circadian phenotypes ^67–69^. While the number of candidate loci has surged in recent years, these results have not yet been translated to biological insights into sleep/wake regulation or the pathophysiology of sleep and circadian disorders. These GWAS data have also demonstrated significant pleiotropy, with loci associated with both sleep phenotypes and mental health traits in particular. Identifying causal genes for sleep-related traits may thus also yield insights into the genetic architecture of psychiatric disorders. With the growth of biobanks in multiple health systems, there are unique opportunities to identify individuals with common and rare genetics variants in *PIG-Q*, and other causal genes, for in-depth sleep and psychiatric phenotyping to improve our understanding of the potential functional effects of these genes in humans.

## METHODS

### Genome-Wide Association Study of Insomnia

Summary statistics from the insomnia GWAS meta-analysis published by Jansen et al (combined sample size of 1,331,010 participants) were used to identify our initial pool of candidate variants. The meta-analysis identified ∼12,000 genome-wide significant variants (P < 5 × 10^−8^), located in 202 genomic risk loci. In the UK Biobank^8^.

### Cell culture

Frozen NPCs derived from iPSC from two healthy individuals (WT10 and WT14) were obtained from CHOP stem cell core and thawed slowly in 37°C water bath. The thawed cells were gently washed in Neuronal Expansion Media: 49% Neurobasal Media (ThermoFisher, cat# 21103049), 49% Advanced DMEM/F12 (ThermoFisher, cat# 12634010) and 2% 50X Neuronal Induction Supplement) in a 15 mL conical tube, followed by centrifuging at 300x g for 5 mins. Cells were resuspended in 1 mL pre-warmed Neuronal Expansion Media with Rock inhibitor (Y-27632 compound, Stem Cell Technologies, cat# 72304) at a final concentration of 10 μM and a cell count performed. NPCs were seeded at a density of 150k cells/cm2 onto hESC-qualified Matrigel-coated plates (Corning, cat# 354277) in 2.5 mL/well Neuronal Expansion Media and cultured at 37°C in a humidified cell culture incubator with 5% CO2. The day after, the medium was changed to remove the Y-27632 compound. NPCs were expanded for 6-7 days in 2.5 mL Neuronal Expansion Media exchanged every 48 hours before harvesting.

### ATAC-seq library preparation

Five technical replicates of two iPSC-derived NPC lines (CHOPWT10 and CHOPWT14) were harvested using Accutase, followed by a DPBS wash, then counted. 50,000 cells of each sample were spun down at 550 ×g for 5 min at 4°C. The cell pellet was then resuspended in 50μL cold lysis buffer (10 mM Tris-HCl, pH 7.4, 10 mM NaCl, 3 mM MgCl2, 0.1% IGEPAL CA-630) and spun down immediately at 550 ×g for 10 min, 4°C. The nuclei were resuspended on ice in the transposition reaction mix (2x TD Buffer, 2.5μL Tn5 Transposes and Nuclease Free H_2_O) (Illumina Cat #FC-121-1030, Nextera) on ice and the transposition reaction was incubated at 37°C for 45 min. The transposed DNA was then purified using a MinElute Kit (Qiagen) adjusted to 10.5 μl elution buffer. The transposed DNA was converted into libraries using NEBNext High Fidelity 2x PCR Master Mix (NEB) and the Nextera Index Kit (Illumina) by PCR amplification for 12 cycles. The PCR reaction was subsequently cleaned up using AMPureXP beads (Agencourt), checked on a Bioanalyzer 2100 (Agilent) high sensitivity DNA Chip (Aglient), and paired-end sequenced on the Illumina NovaSeq 6000 platform (51bp read length) at the Center for Spatial and Functional Genomics at CHOP.

### RNA-seq library preparation

RNA was isolated from two iPSC-derived NPC lines (CHOPWT10 and CHOPWT14) in technical triplicates using Trizol Reagent (Invitrogen). RNA was then purified using the Directzol RNA Miniprep Kit (Zymol) and depleted of contaminating genomic DNA using DNAse I. Purified RNA was then checked for quality on the Bioanlyzer 2100 using the Nano RNA Chip and samples with a RIN number above 7 were used for RNA-seq library synthesis. RNA samples were depleted of rRNA using the QIAseq FastSelect RNA Removal Kit then processed into libraries using the NEBNext Ultra Directional RNA Library Prep Kit for Illumina (NEB) according to manufacturer’s instructions. Quality and quantity of the libraries was measured using the Bioanalyzer 2100 DNA-1000 chip and Qubit fluorometer (Life Technologies). Completed libraries were pooled and sequenced on the NovaSeq 6000 platform using paired-end 51bp reads at the Center for Spatial and Functional Genomics at CHOP.

### Promoter focused Capture-C library preparation

We used standard methods for generation of 3C libraries^14^. For each library, 10^7^ fixed cells were thawed at room temperature, followed by centrifugation at RT for 5 mins at 14,000rpm. The cell pellet was resuspended in 1 mL of dH2O supplemented with 5 μL 200X protease inhibitor cocktail, incubated on ice for 10 mins, then centrifuged. The cell pellet was resuspended to a total volume of 650μL in dH2O. 50μL of cell suspension was set aside for pre-digestion QC, and the remaining sample was divided into 6 tubes. Both pre-digestion controls and samples underwent a pre-digestion incubation in a Thermomixer (BenchMark) with the addition of 0.3%SDS, 1x NEB DpnII restriction buffer, and dH2O for 1hr at 37°C shaking at 1,000rpm. A 1.7% solution of Triton X-100 was added to each tube and shaking was continued for another hour. After the pre-digestion incubation, 10 μl of DpnII (NEB, 50 U/µL) was added to each sample tube only, and continued shaking along with pre-digestion control until the end of the day. An additional 10 µL of DpnII was added to each digestion reaction and digested overnight. The next day, a further 10 µL DpnII was added and continue shaking for another 2-3 hours. 100 μL of each digestion reaction was then removed, pooled into two 1.5 mL tube, and set aside for digestion efficiency QC. The remaining samples were heat inactivated at 1000 rpm in a MultiTherm for 20 min, at 65°C, and cooled on ice for 20 minutes. Digested samples were ligated with 8 μL of T4 DNA ligase (HC ThermoFisher, 30 U/µL). and 1X ligase buffer at 1,000 rpm overnight at 16°C in a MultiTherm. The next day, an additional 2 µL of T4 DNA ligase was spiked into each sample and incubated for another few hours. The ligated samples were then de-crosslinked overnight at 65°C with Proteinase K (20 mg/mL, Denville Scientific) along with pre-digestion and digestion control. The following morning, both controls and ligated samples were incubated for 30 min at 37°C with RNase A (Millipore), followed by phenol/chloroform extraction, ethanol precipitation at -20°C, then the 3C libraries were centrifuged at 3000 rpm for 45 min at 4°C to pellet the samples. The controls were centrifuged at 14,000 rpm. The pellets were resuspended in 70% ethanol and centrifuged as described above. The pellets of 3C libraries and controls were resuspended in 300μL and 20μL dH2O, respectively, and stored at −20°C. Sample concentrations were measured by Qubit. Digestion and ligation efficiencies were assessed by gel electrophoresis on a 0.9% agarose gel and by quantitative PCR (SYBR green, Thermo Fisher).

Isolated DNA from 3C libraries was quantified using a Qubit fluorometer (Life technologies), and 10 μg of each library was sheared in dH2O using a QSonica Q800R to an average fragment size of 350bp. QSonica settings used were 60% amplitude, 30s on, 30s off, 2 min intervals, for a total of 5 intervals at 4 °C. After shearing, DNA was purified using AMPureXP beads (Agencourt). DNA size was assessed on a Bioanalyzer 2100 using a DNA 1000 Chip (Agilent) and DNA concentration was checked via Qubit. SureSelect XT library prep kits (Agilent) were used to repair DNA ends and for adaptor ligation following the manufacturer protocol. Excess adaptors were removed using AMPureXP beads. Size and concentration were checked again by Bioanalyzer 2100 using a DNA 1000 Chip and by Qubit fluorometer before hybridization. One microgram of adaptor-ligated library was used as input for the SureSelect XT capture kit using manufacturer protocol and our custom-designed 41K promoter Capture-C probe set. The quantity and quality of the captured libraries were assessed by Bioanalyzer using a high sensitivity DNA Chip and by Qubit fluorometer. SureSelect XT libraries were then paired-end sequenced on Illumina NovaSeq 6000 platform (51bp read length) at the Center for Spatial and Functional Genomics at CHOP.

### ATAC-seq peak calling

NPC ATAC-seq peaks were called using the ENCODE ATAC-seq pipeline (https://www.encodeproject.org/atac-seq/) with the optimal peaks IDR option.

### Promoter Capture-C pre-processing and interaction calling

Paired-end reads from NPCs were pre-processed using the HICUP pipeline ^37^ (v0.5.9), with bowtie2^78^ as aligner and hg19 as the reference genome. Non-hybrid read count from all baited promoters were used for significant promoter interaction calling. Significant promoter interactions at 1-DpnII fragment resolution were called using CHiCAGO ^79^ (v1.1.8) with default parameters except for binsize set to 2500. Significant interactions at 4-DpnII fragment resolution were also called using CHiCAGO with artificial baitmap and rmap files in which DpnII fragments were concatenated *in silico* into 4 consecutive fragments. Interactions with a CHiCAGO score > 5 in at least one cell type in either 1-fragment or 4-fragment resolution were considered as significant interactions. The significant interactions were finally converted to ibed format in which each line represents a physical interaction between fragments.

### RNA-seq expression analysis

STAR (Spliced Transcripts Alignment to a Reference, v2.5.2b) was used to align each paired-end fastq file for each RNA-seq library independently to reference GRCh37. GencodeV19 was used for gene feature annotation and the raw read count for gene feature was calculated by htseq-count (v0.6.1) with parameter settings -f bam -r pos -s reverse -t exon -m union [20]. The gene features localized on chrM or annotated as rRNAs were removed from the final sample-by-gene read count matrix. Transcript Per Million (TPM) and percentile expression values were calculated from the raw read counts for each gene with a custom script in R, using GencodeV19 annotation for gene lengths.

### Variant to gene mapping

Proxy SNPs for each sentinel SNP reported in Jansen et al. were calculated using online SNP annotator SNiPA (https://snipa.helmholtz-muenchen.de/snipa/) (settings: genome assembly as GRCh37, variant set as 1000 Genome Phase 3 v5, LD r-square cutoff as 0.7) in the European population. Proxy SNPs positions were intersected with the position of the NPC ATAC-seq peaks to identify open “informative” proxies. Capture C chromatin loops to open gene promoters were annotated to each open proxy SNP using custom scripts.

### *Drosophila* husbandry

Flies were grown and maintained on standard *Drosophila* food media (Bloomington Recipe, Genesee Scientific, San Diego, California) in incubators (Powers Scientific, Warminster, Pennsylvania) at 25°C on a 12:12 LD cycle with humidity set to 55–65%. The following fly strains were obtained from the Bloomington Stock Center (Ni et al., 2009): *w*^1118^ (#5905), nsyb-GAL4 (#39171), UAS-*PIG-Q* RNAi^1^ (#67955), while the UAS-*PIG-Q* RNAi^2^ (#107774) was obtained from the Vienna *Drosophila* Resource Center ^70^ or the Bloomington Stock Center ^71^. The stock numbers of all lines used for screeding are described in Supplemental Tables 2-4 unless otherwise stated. Mated females aged 3-to-5 days were used for all experiments performed in this study._7071_

### Sleep and Arousal Threshold Measurements

Flies were acclimated to experimental conditions for at least 24 hrs prior to the start of all behavioral analysis. Measurements of sleep and arousal threshold were then measured over the course of three days starting at ZT0 using the *Drosophila* Locomotor Activity Monitor (DAM) System (Trikinetics, Waltham, MA, USA), as previously described ^30,72,73^. For each individual fly, the DAM system measures activity by counting the number of infrared beam crossings over time. These activity data were then used to calculate sleep, defined as bouts of immobility of 5 min or more, using the *Drosophila* Sleep Counting Macro ^74^, from which sleep traits were then extracted. Waking activity was quantified as the average number of beam crossings per waking minute, as previously described ^74^.

Arousal threshold was measured using the *Drosophila* Arousal Tracking system (DART), as previously described ^35^. In brief, individual female flies were loaded into plastic tubes (Trikinectics, Waltham, Massachusetts) and placed onto trays containing vibrating motors. Flies were recorded continuously using a USB-webcam (QuickCam Pro 900, Logitech, Lausanne, Switzerland) with a resolution of 960×720 at 5 frames per second. The vibrational stimulus, video tracking parameters, and data analysis were performed using the DART interface developed in MATLAB (MathWorks, Natick, Massachusetts). To track fly movement, raw video flies were subsampled to 1 frame per second. Fly movement, or a difference in pixilation from one frame to the next, was detected by subtracting a background image from the current frame. The background image was generated as the average of 20 randomly selected frames from a given video. Fly activity was measured as movement of greater than 3 mm. Sleep was determined by the absolute location of each fly and was measured as bouts of immobility for 5 min or more. Reactivity was assessed using a vibration intensity of 1.2 g, once per hour over 3 days starting at ZT0.

All measurements of sleep and arousal threshold were combined across the three days of testing. Statistical analyses were performed in Prism (GraphPad Software 9.3). Unless otherwise noted, a t-test or one-way analysis of variance (ANOVA) was used for comparisons between two genotypes or two or more genotypes, respectively. All *post hoc* analyses were performed using Dunnett’s multiple comparisons test. Measurements of arousal threshold were not normally distributed and so the non-parametric restricted maximum likelihood (REML) estimation was used. To characterize the relationship between the change in reactivity and bout length, we performed linear regression analyses. An ANCOVA was used to compare the elevations of different genotypes.

### Sleep Deprivation

Sleep deprivation experiments were performed as previously described (Murakami et al., 2016). Upon experiment onset, baseline sleep was measured starting at ZT0 for 24 hrs. For the following 24 hrs, flies were mechanically sleep deprived, during which sleep was also measured. To assess homeostatic rebound, flies were returned to standard conditions and sleep was measured during the subsequent day (ZT0-ZT12). To determine whether there exists a homeostatic rebound in sleep duration, baseline daytime sleep was compared to daytime sleep during recovery.

### Generation of zebrafish mutant

*PIG-Q* knockout (KO) mutants were generated using CRISPR/Cas9 as previously described ^76^. Single cell-stage embryos were injected with preformed RNP complexes containing Cas9 protein and an sgRNA, which has a target sequence of 5’**-**TCTAAAGAGTCGCCAGAGCGAGG-3’. Scrambled sgRNA with sequence 5’-CGTTAATCGCGTATAATACG-3’ was used for negative control injections. Mutant animals were genotyped using Headloop PCR as described in ^43^, and larvae were included in sleep assay analysis if they had a mutation efficiency greater than 0.9 (90%) as determined by the ratio of Headloop to standard PCR product using ImageJ. Primers for the target region included standard primers 5’-GTTGGAGTGACTCACCAGGG-3’ and 5’-TGAGTACTGCAGGGTGGTTTC-3’ and Headloop primers 5’-GTTGGAGTGACTCACCAGGG-3’ and 5’-AGAGCGAGGAGAGACCGTAGTGAGTACTGCAGGGTGGTTTC-3’. Guide RNA design was performed using CRISPOR tefor (http://crispor.tefor.net/) to optimize sensitivity and specificity >95%, with minimal off-target effects. CRISPR sgRNA is designed for the exonic region with highest conservation across species using MARRVEL ^44^ and is a component of the N-acetylglucosaminyl transferase component that overlaps between both *PIG-Q* transcripts to ensure disruption of all possible transcripts. Target region was also free of *in silico* predicted SNPs and Sanger sequencing was performed to experimentally demonstrate target region was free of mutations.

### Sleep/wake assay in zebrafish

Zebrafish experiments were performed in accordance with University of Pennsylvania Institutional Animal Care and Use Committee (IACUC) guidelines (animal protocol # 806646). Zebrafish embryos were collected from AB/TL incross breeding pairs on the morning of spawning and injected with CRISPR/Cas9 reagents at the single cell stage. Embryos were raised on a 14:10 hour light:dark cycle. Animals were housed in petri dishes with approximately 50 per dish in standard embryo medium (E3 medium; 5mM NaCl, 0.17 mM KCl, 0.33 mM CaCl2, 0.33 mM MgSO_4,_ 10^−5^ % Methylene Blue) and kept in an incubator at 28.5°C. Dead embryos and shed chorion membranes were removed on days 2-4 post-fertilization. Scramble guide RNA-injected and *PIG-Q* knockout larvae were individually placed into each well in alternating rows of a 96-well plate in 650 μL of E3 embryo medium without Methylene Blue on 5 days post-fertilization. Activity was captured using automated video tracking (Viewpoint Life Sciences) for 72 hours. Behavioral phenotyping of larvae at F0 generation, which display high mutation efficiencies greater than 90 percent has been modified from ^43^. The 96-well plate was housed in a Zebrabox (Viewpoint Life Sciences) with customizable light parameters and a Dinion one-third inch Monochrome camera (Dragonfly2, Point Grey) fitted with a variable-focus megapixel lens (M5018-MP, Computar) and infrared filter. The plate was fitted in a chamber filled with recirculating water connected to a temperature-control unit to maintain a stable temperature of 28.5°C which is the optimum growth temperature of zebrafish. Activity was captured in quantization mode with detection parameters: threshold, 20; burst, 29; freeze, 3; bin size, 60 sec as described previously ^77^.

### Zebrafish sleep/wake data analysis

An acclimation period was removed, and data analysis consisted of two days and two nights. Data were processed using custom MATLAB (The Mathworks, Inc) scripts as performed in^77^ with modifications. Movement was captured as seconds per minute and any one-minute period with less than 0.5 sec of total movement was defined as one minute of sleep (modified from ^78^). Sleep bouts were defined as a continuous string of sleep defined in minutes and sleep bout length was calculated as minutes across the continuous string. Average activity was defined as the average amount of activity using the threshold of 0.5 sec to define waking activity and reported as seconds per awake minute per hour. Statistical tests were performed in Prism (Graphpad). Activity was combined across both days and sleep across both nights for analysis. Independent student’s t-test was used to compare groups with equal variances, while Welch’s correction was applied when variances were determined to be unequal.

### DNA Extraction and PCR Validation

DNA extraction was performed per the manufacturer’s protocol (Quanta bio, Beverly, MA) immediately following completion of the sleep assay. Larvae were euthanized by rapid cooling on a mixture of ice and water between 2-4°C for a minimum of 30 minutes after complete cessation of movement was observed. Larvae were transferred to a new 96-well PCR plate and excess E3 buffer was removed. Twenty-five μL of DNA extraction buffer was then added and larvae were completely submerged. The plate was sealed and heated for 30 minutes at 95°C then cooled to room temperature. Twenty-five μL of DNA stabilization buffer was then added and genomic DNA was stored at 4°C. For PCR validation, each well of a PCR plate contained 0.1 μL of Phusion DNA Polymerase, 5 μL of 5x Phusion HF Buffer, 0.5 μL of dNTP mix, 0.5 μL of 10 μM *PIG-Q* forward primer and 0.5 μL of 10 μM *PIG-Q* reverse primer or headloop antisense primer, 16.4 μL of nuclease free water and 2 μL of two-fold diluted genomic DNA for a final volume of 25 μL. PCR plate was sealed and placed into a thermocycler. The PCR reaction conditions were: one cycle of 98°C for 90 sec; 30 cycles of 98°C for 10 sec, 64°C for 10 sec, 72°C for 15 sec, 72°C for 15 sec; one cycle of 72°C for 5 min, then stored at 4°C. Samples were run on 2% agarose gel and quantified using ImageJ for the ratio of headloop PCR product to standard PCR product to calculate mutation efficiency as previously described ^43^.

## ACKNOWLEDGEMENTS

This work was supported by NIH grant 5R01HL143790 to ACK, SFAG, and PG, award 1R21NS122166 to ACK, and K99AB071833 to EBB. SFAG is also supported by NIH grants R01 HD056465, R01 AG057516, and the Daniel B. Burke Endowed Chair for Diabetes Research. AC was supported by NIH grant R35 HG011959. AIP was supported by P01 HL094307, and T32 HL07953.

## Supplementary Tables

**Table S1. Candidate target genes identified for the insomnia GWAS loci by our variant-to-gene mapping approach**. GWAS loci are numbered according to ^19^ Candidate regulatory proxy SNPs are reported with the r^2^ value (in EUR population) to their respective sentinel SNPs. Candidate target genes are reported with their TPM (transcript per million) expression and percentile expression values from our bulk RNA-seq data in neural progenitor cells.

**Table S2. RNAi screen of insomnia-associated orthologs**. Raw sleep data from sleep screen includes stock numbers of each RNAi line used as well as the mean measurements of the other measurements of sleep including sleep duration, bout number, average bout length and waking activity.

**Table S3. Localization of PIG-Q-RNAi long sleeping phenotype**. Raw sleep data after knockdown of *PIG-Q* in multiple *Drosophila* neuronal subpopulations.

**Table S4. Pan-neuronal knockdown of PIG pathway genes**. Raw sleep data after pan-neuronal knockdown of PIG pathway genes.

## Supplementary Figures

**Figure S1.**
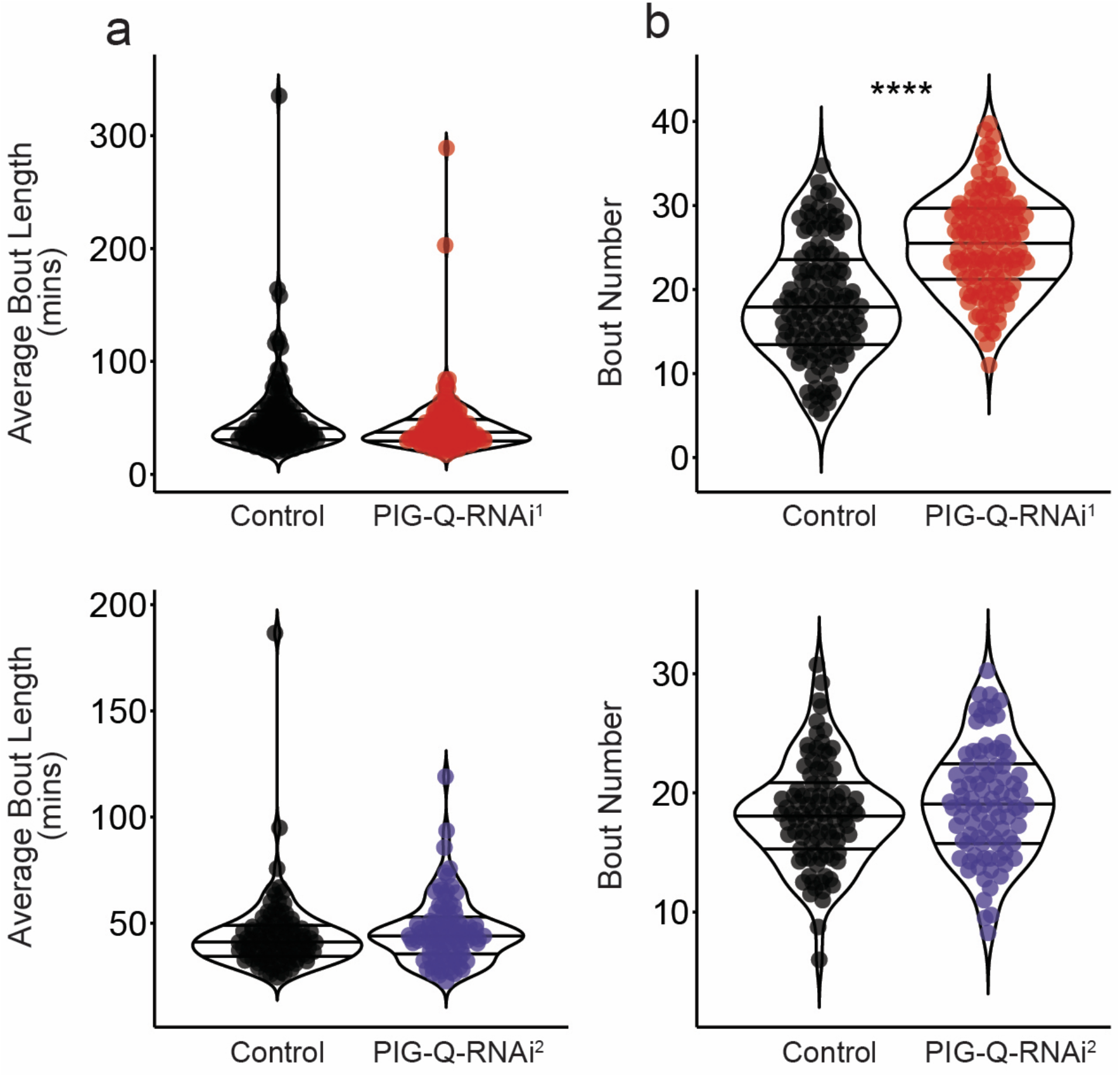
The effect of *PIG-Q* pan-neuronal knockdown on sleep traits. **a**, Knockdown of *PIG-Q* has no effect on bout length in two independent RNAi lines (t-test, PIG-Q-RNAi^1^: t_260_ = 1.668; *P*=0.0966; PIG-Q-RNAi^2^: t_184_ = 0.8104; *P*=0.4188). **b**, Knockdown of *PIG-Q* significantly increases bout number in one RNAi line, but has no effect in a second independent RNAi line (t-test, *PIG-Q*-RNAi^1^: t_260_ = 8.949; *P*<0.0001; PIG-Q-RNAi^2^: t_184_ = 1.470; *P* = 0.1432). \*\*\*\**P*<0.0001.

**Figure S2.**
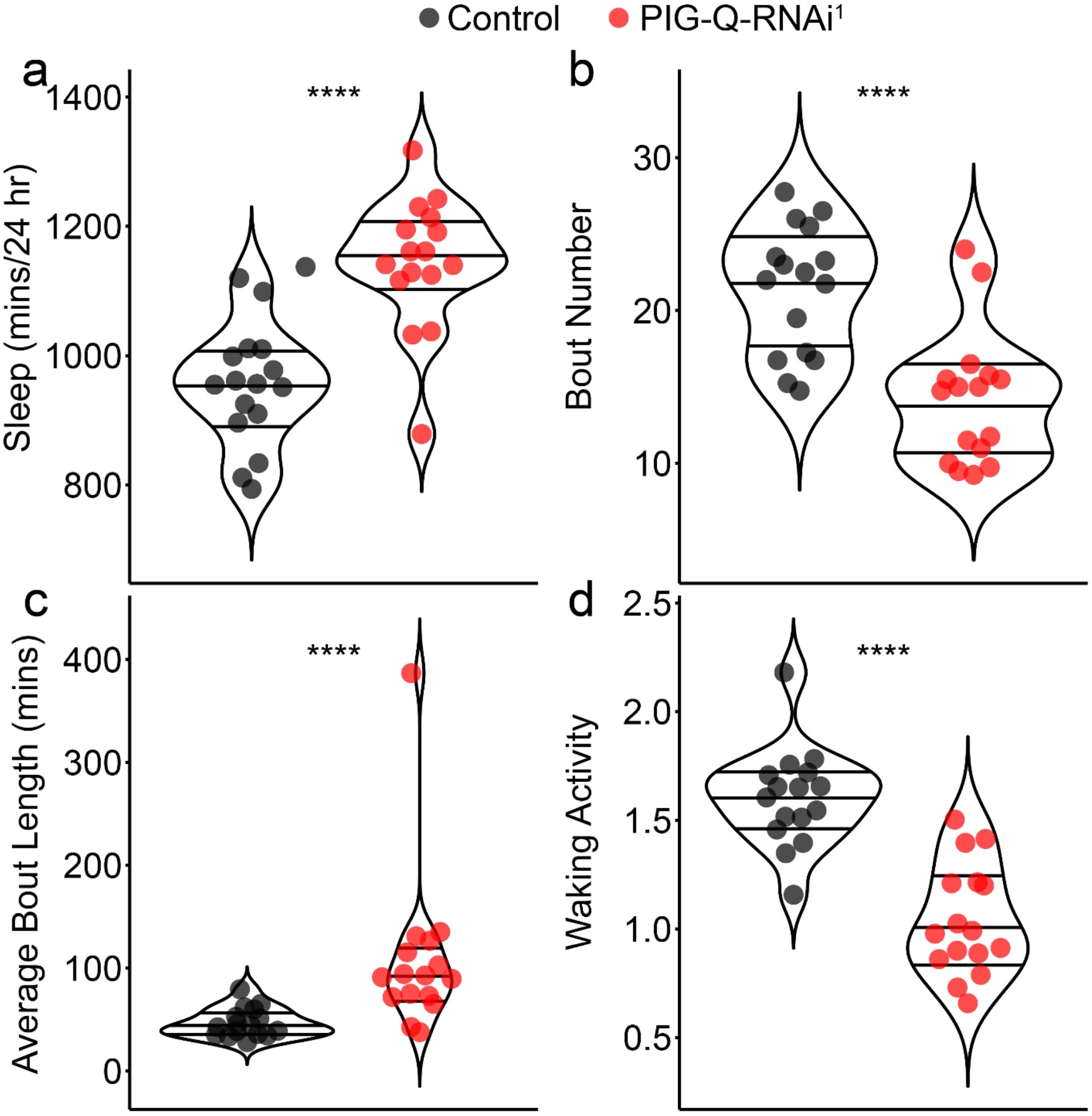
The effect of *PIG-Q* pan-neuronal knockdown on sleep traits in male flies. **a**, Knockdown of *PIG-Q* significantly increases sleep duration (t-test, t_30_ = 5.708; *P*<0.0001). **b**, Knockdown of *PIG-Q* significantly increases the length of each sleep episode (t-test, t_30_ = 3.042; *P*<0.0001), while significantly decreasing the number of sleep bouts (**c**, t-test, t_30_ = 4.748; *P*<0.0001). **d**, Knockdown of *PIG-Q* significantly decreases waking activity (t-test, t_30_ = 6.596; *P*<0.0001). \*\*\*\**P*<0.0001.

**Figure S3.**
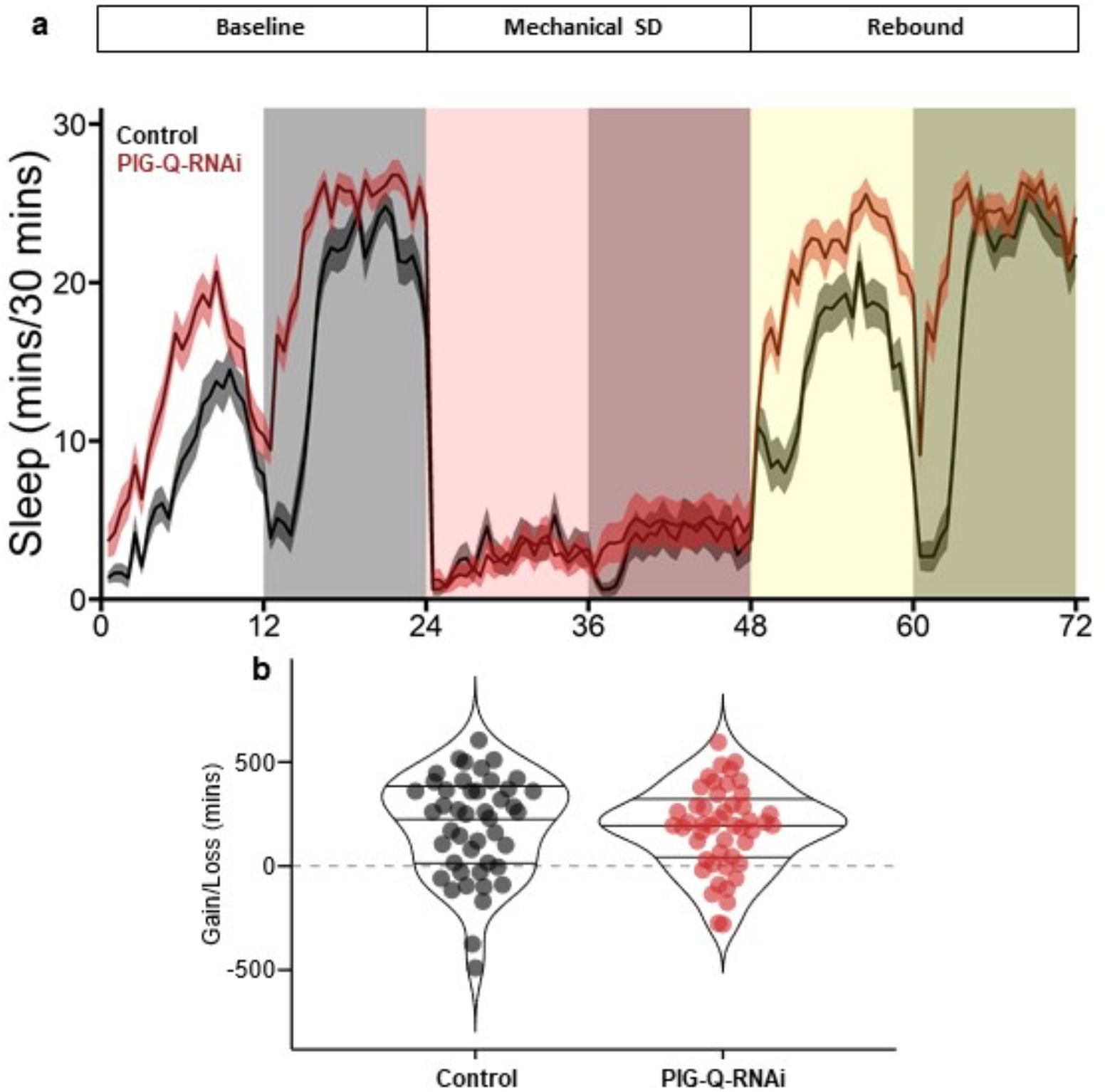
*PIG-Q* pan-neuronal knockdown has no effect on homeostatic sleep. **a**, Sleep profile of flies before (white), during (red), and after (yellow) 24 hrs of mechanical sleep deprivation. **b**, Knockdown of *PIG-Q* has no effect on sleep rebound. Minutes gained/lost are determined by measuring the acclimation period (white) and subtracting it from the rebound period (yellow) (t-test; t_55_ = 7.387; *P*=0.4816).

**Figure S4.**
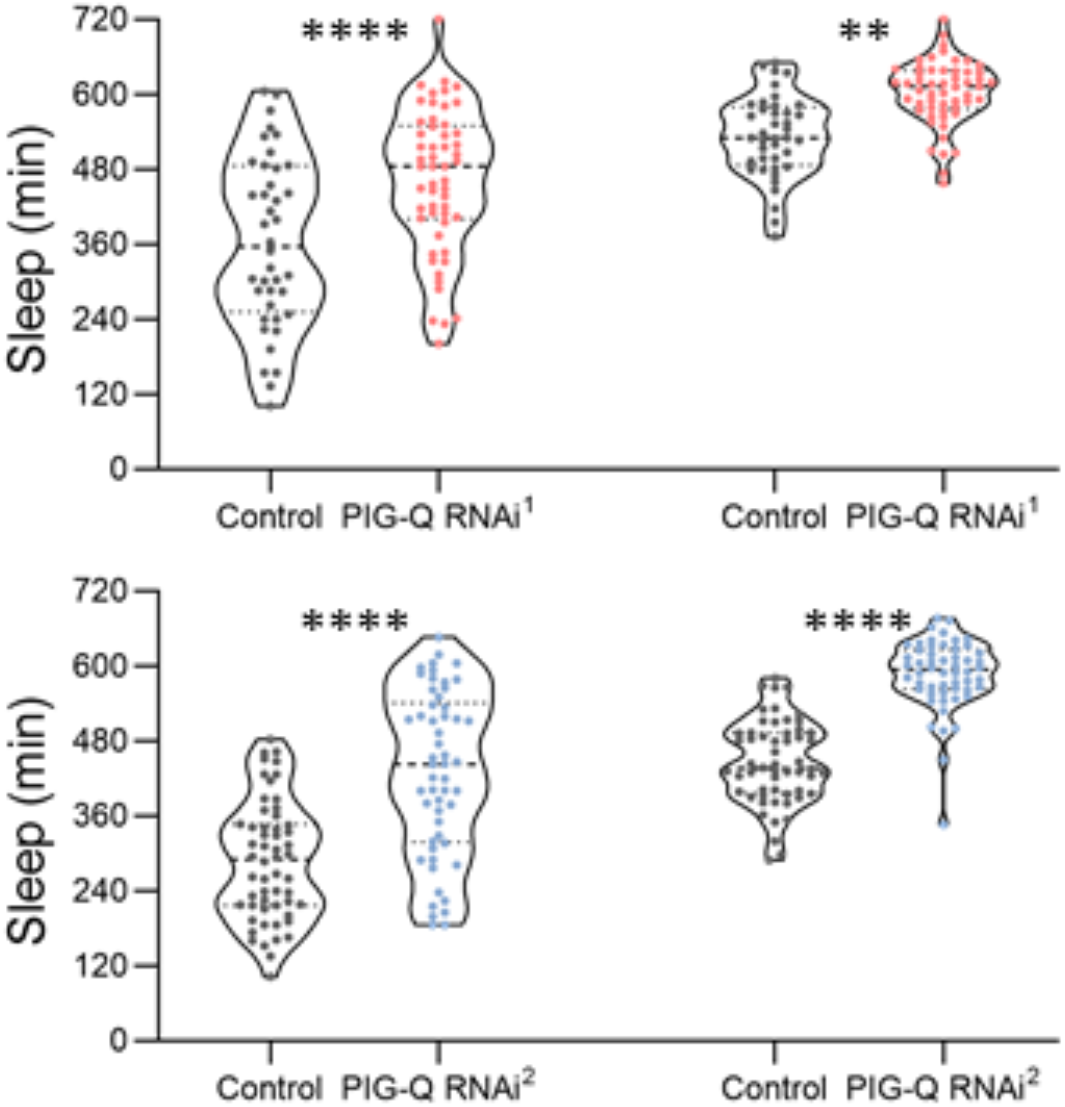
Sleep duration measurements of *PIG-Q* pan-neuronal knockdown in the DART system. **a**, Knockdown of PIG-Q significantly increases sleep in two independent RNAi lines (*PIG-Q*-RNAi^1^: F_1,184_ = 34.75, *P*<0.0001; *PIG-Q*-RNAi^2^: F_1,148_ = 37.71, *P*<0.0001), and occurs during both the day (*PIG-Q*-RNAi^1^: *P*<0.0001; *PIG-Q*-RNAi^2^: *P*<0.0001) and night (*PIG-Q*-RNAi^1^: *P*=0.0014; *PIG-Q*-RNAi^2^: *P*<0.0001). \*\**P* < 0.01, \*\*\*\**P* < 0.0001.

**Figure S5.**
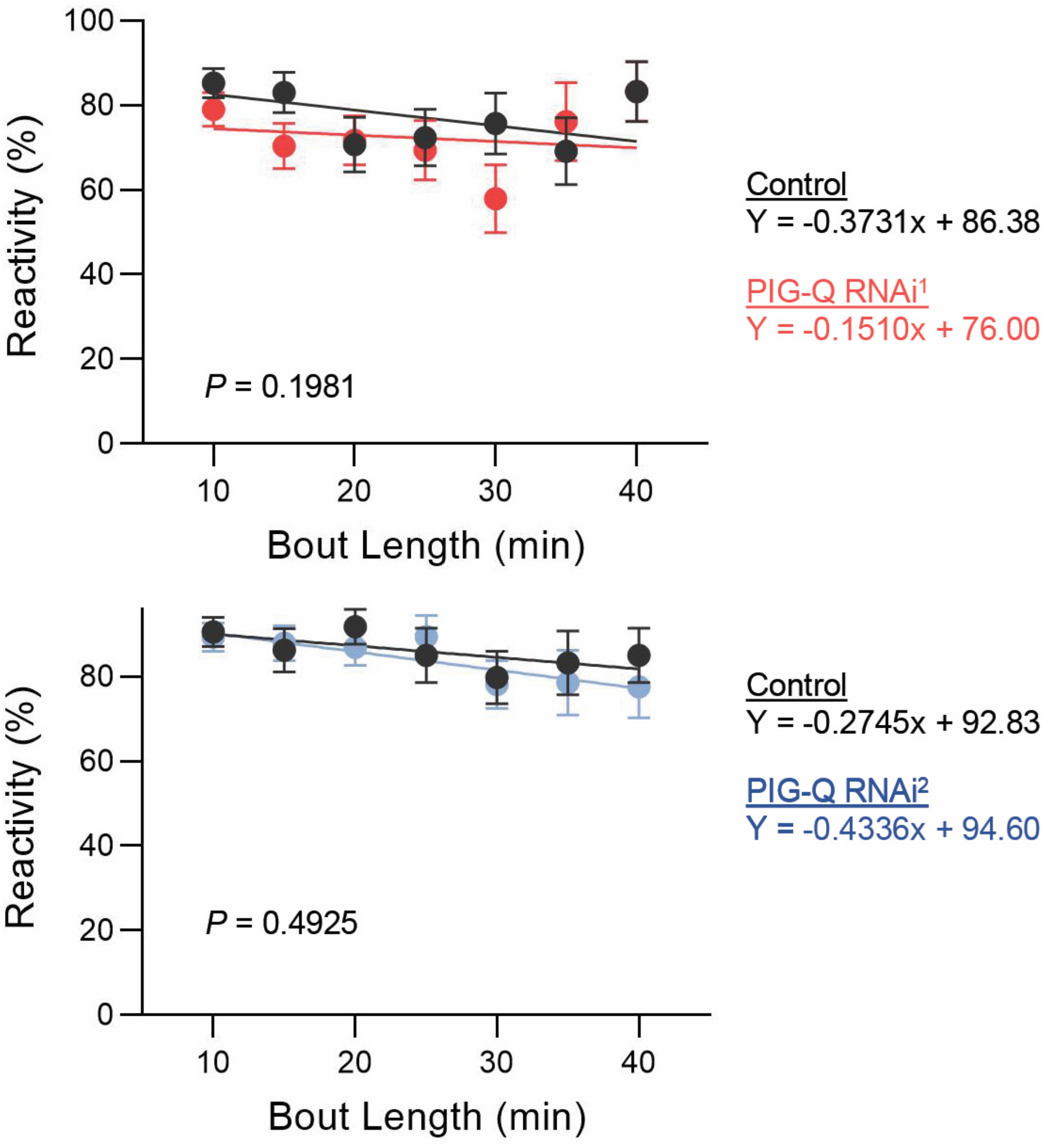
Daytime reactivity of pan-neuronal knockdown of *PIG-Q*. Knockdown of *PIG-Q* has no effect on daytime reactivity (ANCOVA with bout length as covariate, *PIG-Q*-RNAi^1^: F_1,715_ = 0.4631, *P*=0.4964; *PIG-Q*-RNAi^2^: F_1,637_ = 0.3389, *P*=0.5606).

